# Engineered shell proteins confer improved encapsulated pathway behavior in a bacterial microcompartment

**DOI:** 10.1101/106716

**Authors:** Marilyn F. Slininger Lee, Christopher M. Jakobson, Danielle Tullman-Ercek

## Abstract

Bacterial microcompartments are a class of proteinaceous organelles comprising a characteristic protein shell enclosing a set of enzymes. Compartmentalization can prevent escape of volatile or toxic intermediates, prevent off-pathway reactions, and create private cofactor pools. Encapsulation in synthetic microcompartment organelles will enhance the function of heterologous pathways, but to do so, it is critical to understand how to control diffusion in and out of the microcompartment organelle. To this end, we explored how small differences in the shell protein structure result in changes in the diffusion of metabolites through the shell. We found that the ethanolamine utilization (Eut) protein EutM properly incorporates into the 1,2-propanediol utilization (Pdu) microcompartment, altering native metabolite accumulation and the resulting growth on 1,2-propanediol as the sole carbon source. Further, we identified a single pore-lining residue mutation that confers the same phenotype as substitution of the full EutM protein, indicating that small molecule diffusion through the shell is the cause of growth enhancement. Finally, we show that the hydropathy index and charge of pore amino acids are important indicators to predict how pore mutations will affect growth on 1,2- propanediol, likely by controlling diffusion of one or more metabolites. This study highlights the success of two strategies to engineer microcompartment control over metabolite transport: altering the existing shell protein pore via mutation of the pore-lining residues, and generating chimeras using shell proteins with the desired pores.

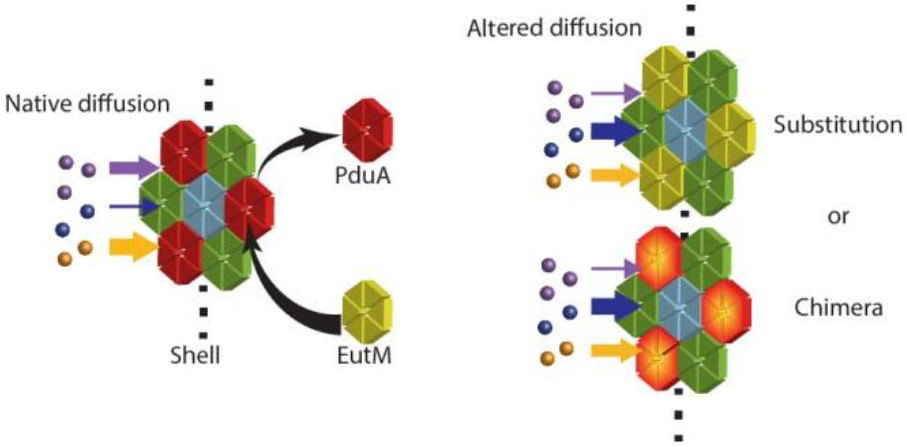
TOC Abstract Graphic.

---

Bacterial microcompartments (MCPs) are a prokaryotic form of cellular organization consisting of a protein shell used to enhance pathways by encapsulation. MCP-associated pathways include those found in hosts across many bacterial phyla for metabolism of niche carbon sources, such as 1,2-propanediol (1,2-PD), ethanolamine, and ethanol (involving Pdu, Eut, and Etu MCPs, respectively), and those found in cyanobacteria and chemoautotrophs for carbon fixation by RuBisCo in the carboxysome.^1#4^ Additional naturally-encapsulated pathways have been recently discovered and a summary of this topic may be found in a number of review articles.^5#8^ Some bacterial species have several MCP systems, encoding different organelles tailored to metabolize specific substrates.^9#10,11,12^ For example, two well-characterized model systems are found in *Salmonella enterica,* which can produce both the Eut and Pdu MCPs from two separate operons. These enable growth on ethanolamine and 1,2-PD, respectively, which are both found in the human gut. The Pdu MCP encapsulates enzymes converting 1,2-PD through a propionaldehyde intermediate into either 1-propanol or propionate (Figure 1a), while the Eut MCP similarly converts ethanolamine through an acetaldehyde intermediate into ethanol and acetyl phosphate. The benefits of encapsulating a pathway include preventing the escape or toxicity of intermediates as well as cycling of cofactors.^13#17^

The *S. enterica* Pdu MCP assembly is an irregular polyhedron 150-200 nm in diameter and is enclosed by a shell composed of a few thousand copies of shell proteins of several types.^18^ There are eight Pdu shell protein paralogs that are present in the final assembly in distinct and reproducible ratios.^19^ The *S. enterica* Eut MCP is similarly composed of five shell proteins. MCP shell proteins form trimers, pentamers, or hexamers that come together to assemble the hexagonal or pentagonal tiles of the polyhedron. A pore is formed at the symmetric center of each tile.^18–21^

The central pores of MCP shell proteins are implicated in the transport of small molecule substrates and products across the shell.^22, 23^ For instance, Pdu MCP shell protein PduA is abundant in the shell and is suggested to be involved in the selective transport of 1,2-PD.^23^ Specifically, occlusion of its pore by PduA^S40GSG^, PduA^S40L^, PduA^S40C^, and PduA^S40Q^ mutations lead to growth defects that are rescued by high 1,2-PD concentrations, while the PduA^S40A^ mutation increased the escape of propionaldehyde.^23^ Similarly, the pores of the carboxysome are hypothesized to control flux of bicarbonate and CO2 across the shell.^24^ All characterized carboxysome pore structures are positively charged at the narrowest point, which could enhance the flux of negatively charged bicarbonate relative to the escape of CO2. Since different pore structures are postulated to lead to altered diffusivity of metabolites, engineering the pores could be the key to tailoring MCP properties for encapsulating non-native systems.

An alternative strategy for engineering MCP properties for non-native pathways is to use novel combinations of natural shell proteins. Shell proteins from different MCP systems have the potential to interact to form chimeric assemblies if expressed together at appropriate levels. Cai et al. demonstrated that an alpha-carboxysome shell protein complements the deletion of a beta-carboxysome shell protein.^25^ Sturms et al. observed that co-expression of the Eut and Pdu operons in *S. enterica* from a synthetic promoter system resulted in co-purification of Eut and Pdu components, albeit with a disruption of Pdu MCP function.^26^ Such observations are consistent with another finding: among many shell protein homologs, there are essential asparagine, arginine and lysine residues at the edge of each tile that are important for assembly, indicating a common interaction mechanism (For PduA these residues are K26, N29, and R79).^4, 27^ In addition, shell assembly does not always depend on cargo loading. The protein shell can form when no cargo is expressed, and foreign proteins may also be encapsulated, or localized to the interior of the shell.^28#31^ In fact, peptide sequences that direct native enzymes to the interior of the Eut MCP also function to encapsulate proteins within the Pdu MCP.^32^ Thus, it may be possible to bring Pdu and Eut shell proteins together to form chimeric MCP assemblies with new properties.

In this work, we explore two strategies for altering MCP function: mutations to the pore residues and generation of chimeric MCPs. We study the well-characterized *S. enterica* Pdu MCP, with a focus on its PduA shell protein and its paralog EutM, also from *S. enterica*. We find that EutM behaves as a PduA substitute in the Pdu MCP, and we describe the phenotypic changes resulting from EutM substitution. A *pduA: eutM* chromosomally modified strain incorporates EutM into the Pdu MCP in place of PduA. Interestingly, the *pduA: eutM* strain exhibits improved growth on 1,2-PD over cells expressing the native Pdu MCP, and this new phenotype is recapitulated by a single amino acid mutation in the PduA pore, PduA^K37Q^. A saturation mutagenesis of position 37 of PduA reveals that the pore is highly amenable to mutation, and that the hydrophobicity and charge of the amino acid at this locus are the most important factors affecting growth on 1,2-PD. This study thus demonstrates the potential for success in altering metabolite transport properties with both MCP engineering strategies, enabling MCP design and optimization for synthetic biology applications.

## RESULTS AND DISCUSSION

### The pore of PduA is different from the pore of EutM

PduA forms one of three canonical shell protein types: a hexameric tile complex containing a central pore approximately 6 Å in diameter.^22^ The residues defining the shape and properties of the pore include those contained in a loop at the hexamer symmetry axis with sequence Lys-Ile-Gly-Ser-Gly-Leu. The importance of the K37 and S40 loci of this sequence was previously demonstrated; certain mutations to either of these residues result in diminished growth of *S. enterica* LT2 on 1,2-PD as a sole carbon source.^23, 27^ EutM, a close PduA homolog from the Eut operon, also forms a hexamer with nearly identical predicted structure (RMSD EutM/PduA = 1.004 Å, PDB ID 3MPY (EutM) and 3NGK (PduA)) and a highly similar amino acid sequence (67% identity).^20, 22^ Refer to the protein sequence alignment for PduA and EutM (Figure S1). Key differences between PduA and EutM are found in the pore, particularly at the residues aligning with PduA K37 and S40. The EutM pore loop has the sequence Gln-Ile-Gly-Gly-Gly-Leu (Figure 1b, c). In Figure 1c, the pore-lining lysine residue in PduA and the glutamine in EutM are highlighted in green.

**Figure 1.**
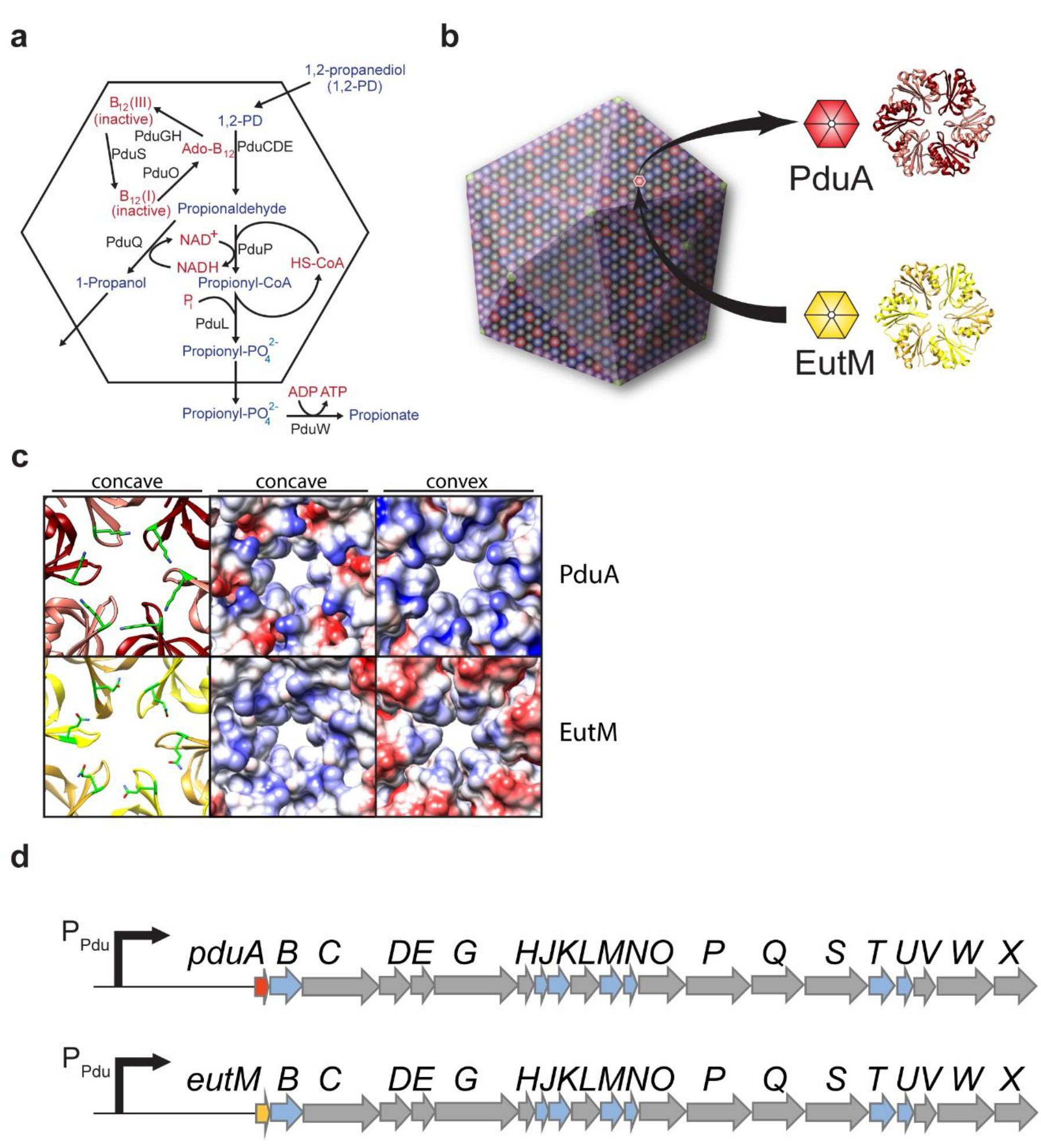
Organization and modifications of Pdu MCP (a)Diagram illustrating the Pdu pathway including enzymes, cofactors, substrates, intermediates and products. The black hexagonal outline represents the boundary of the MCP. Enzymes outside of this boundary are not localized to the interior of the MCP.^39^ (b) Schematic demonstrating the hypothesized overall structure of the MCP. Arrows indicate replacement of the PduA shell protein (red) with EutM (yellow). Crystal structures (PDB IDs: 3ngk, 3mpy) to the right illustrate structural similarity.^14,17^ (c) Left hand images are ribbon structures of the pores of PduA (red) and EutM (yellow). Residues PduA K37 and the corresponding glutamine of the EutM pore are highlighted in green. To the right are the surface models of the respective pore regions colored by the electrostatic potential. Blue is positive potential, white is neutral, and red is negative potential. (d) A diagram of the Pdu operon and the operon resulting from recombineering to substitute eutM for *pduA* on the *S. enterica* chromosome. Other Pdu shell proteins are blue arrows, enzymes of the pathway and proteins of unknown function are grey arrows.

### The growth of *S. enterica* on 1,2PD is improved in*pduA: eutM* strain

We substituted *eutM* for *pduA* to test whether the structural similarity of *eutM* is sufficient to complement the deletion of *pduA* and maintain proper Pdu MCP assembly. We also wanted to learn if small differences between the pore structures would manifest in significant differences in phenotypes attributed to metabolite diffusion. In order to minimize disruption of the native regulation of the Pdu operon, we constructed a genomic integration of *eutM* at the *pduA* locus (Figure 1d). To test MCP formation in this *pduA: eutM* strain, the expression of the Pdu operon was induced in minimal NCE media containing 55 mM 1,2-PD as the sole carbon source, and 150 nM coenzyme B12. Growth of the *pduA: eutM* strain in this medium is an indicator of metabolite diffusion through the MCP structure. For instance, a strain harboring a *pduA* deletion without a compensating substitution exhibits decreased cell growth due to increased shell permeability to the toxic intermediate propionaldehyde, despite producing a seemingly closed MCP that may be purified and appears to contain all other necessary shell proteins and enzymes.^21, 27^ Furthermore, occlusion or other changes to the architecture of the pores of the MCP shell also lead to growth defects due to restriction of diffusion of the substrate 1,2-PD.^7^ Our results show this is not the case for the *pduA: eutM* strain. Not only did the strain harboring the substitution grow without a noticeable defect on 1,2-PD, we observed improved growth over wild type under these growth conditions (Figure 3, Table 1). To control for changes in cell morphology, we also quantified colony forming units at various growth time points to confirm improved growth of the *pduA: eutM* strain (Figure S2). The *pduA: cat/sacb* mutant strain was used as a control for both growth measurements. This strain grew poorly compared to the wild-type (Figure 3, Figure S2). Moreover, no gross changes to cell morphology or physiology were observed in the *pduA: eutM* strain by phase contrast (Figure S3).

**Table 1.** Doubling times of PduA mutants for growth on 1,2-PD. Both *ΔpduA: eutM* and *pduA*
^
*K37Q*
^ strains are statistically different than the wildtype strain by two tailed T-test. The *ΔpduA: eutM* and *pduA*
^
*K37Q*
^ strains are not significantly different from each other by two tailed T-test. n=7 for all cases.

| Strain | Doubling time (hr) | Standard error |
| --- | --- | --- |
| MFSS044, Wildtype | 7.85 | $\pm 0.89$ |
| MFSS252, $\Delta pduA::eutM$ | 5.73 | $\pm 0.29$ |
| MFSS238, $pduA^{K37Q}$ | 5.75 | $\pm 0.49$ |

**Table 2.** Bacterial strains used in this study

| Strain | Organism | Genotype |
| --- | --- | --- |
| MFSS044 | *S. enterica* serovar Typhimurium | LT2 wild type |
| MFSS263 | *S. enterica* serovar Typhimurium | LT2 $pduA^{K26A}$ |
| MFSS236 | *S. enterica* serovar Typhimurium | LT2 $\Delta pduA::cat/sacB$ |
| MFSS252 | *S. enterica* serovar Typhimurium | LT2 $\Delta pduA::eutM$ |
| MFSS273 | *S. enterica* serovar Typhimurium | LT2 $\Delta pduA::eutM$-FLAG |
| MFSS238 | *S. enterica* serovar Typhimurium | LT2 *pduA*<sup>K37Q</sup> |
| MFSS393 | *S. enterica* serovar Typhimurium | LT2 $\Delta pduA::pduA$<sup>K37Q</sup>-FLAG |
| MFSS392 | *S. enterica* serovar Typhimurium | LT2 $\Delta pduA::pduA$-FLAG |
| MFSS337 | *S. enterica* serovar Typhimurium | LT2 $\Delta pduA::eutM$<sup>K24A</sup> |
| MFSS320 | *S. enterica* serovar Typhimurium | LT2 *pduA*<sup>K37H</sup> |
| MFSS321 | *S. enterica* serovar Typhimurium | LT2 *pduA*<sup>K37C</sup> |
| MFSS322 | *S. enterica* serovar Typhimurium | LT2 *pduA*<sup>K37R</sup> |
| MFSS323 | *S. enterica* serovar Typhimurium | LT2 *pduA*<sup>K37G</sup> |
| MFSS324 | *S. enterica* serovar Typhimurium | LT2 *pduA*<sup>K37P</sup> |
| MFSS325 | *S. enterica* serovar Typhimurium | LT2 *pduA*<sup>K37V</sup> |
| MFSS326 | *S. enterica* serovar Typhimurium | LT2 *pduA*<sup>K37F</sup> |
| MFSS327 | *S. enterica* serovar Typhimurium | LT2 *pduA*<sup>K37T</sup> |
| MFSS328 | *S. enterica* serovar Typhimurium | LT2 *pduA*<sup>K37L</sup> |
| MFSS329 | *S. enterica* serovar Typhimurium | LT2 *pduA*<sup>K37D</sup> |
| MFSS330 | *S. enterica* serovar Typhimurium | LT2 *pduA*<sup>K37E</sup> |
| MFSS331 | *S. enterica* serovar Typhimurium | LT2 *pduA*<sup>K37S</sup> |
| MFSS335 | *S. enterica* serovar Typhimurium | LT2 *pduA*<sup>K37N</sup> |
| MFSS336 | *S. enterica* serovar Typhimurium | LT2 *pduA*<sup>K37W</sup> |
| TUC01<sup>a</sup> | *E. coli* | DH10B *cat/sacB* |

<sup>a</sup>Datta et al.<sup>36</sup>

### EutM is localized to the Pdu MCP in the *S. entericapduA: eutM* strain

We next set out to demonstrate that EutM is incorporated into the shell, and to determine if the formation of the shell was altered by this incorporation. We found that substituting *eutM* for *pduA* allowed the formation of chimeric MCPs that were morphologically similar to the wildtype, as assessed by a number of assays. First, we targeted green fluorescent protein (GFP) for localization within the MCPs produced in each of the variants and analyzed assembly formation *in vivo* using fluorescence microscopy. GFP was expressed as a fusion to the PduD^1-20^ encapsulation tag and an SsrA degradation tag such that only encapsulated GFP remains protected from proteolysis by the ClpXP machinery.^31, 33, 34^ This results in bright puncta by fluorescence microscopy. Fluorescent puncta were observed in both *pduA: eutM* and wild-type strains, confirming MCP formation and protein encapsulation *in vivo* (Figure 2a, e, b, f, and). Automated image analysis by Cell Profiler yielded a count of the number of puncta per cell and indicated that the number of chimeric MCPs per cell is similar to the number of wild-type MCPs per cell Figure S5).^35^ Variability in puncta brightness is often observed between cells as well as between puncta within a cell. While we have not investigated the cause of this feature, it appears consistent across both the wild-type and *pduA: eutM* strains.

We next purified MCPs by sedimentation to compare the morphology of chimeric MCPs to wild-type *in vitro*. Transmission electron microscopy (TEM) analysis showed that MCPs purified from the *pduA: eutM* strain are morphologically similar to wild-type MCPs (Figure 2i, j, and). An analysis of MCP diameter demonstrated that MCPs purified from the *pduA: eutM* strain have a similar mean diameter to the wild-type but with a somewhat wider distribution of sizes (Figure S6).

**Figure 2.**
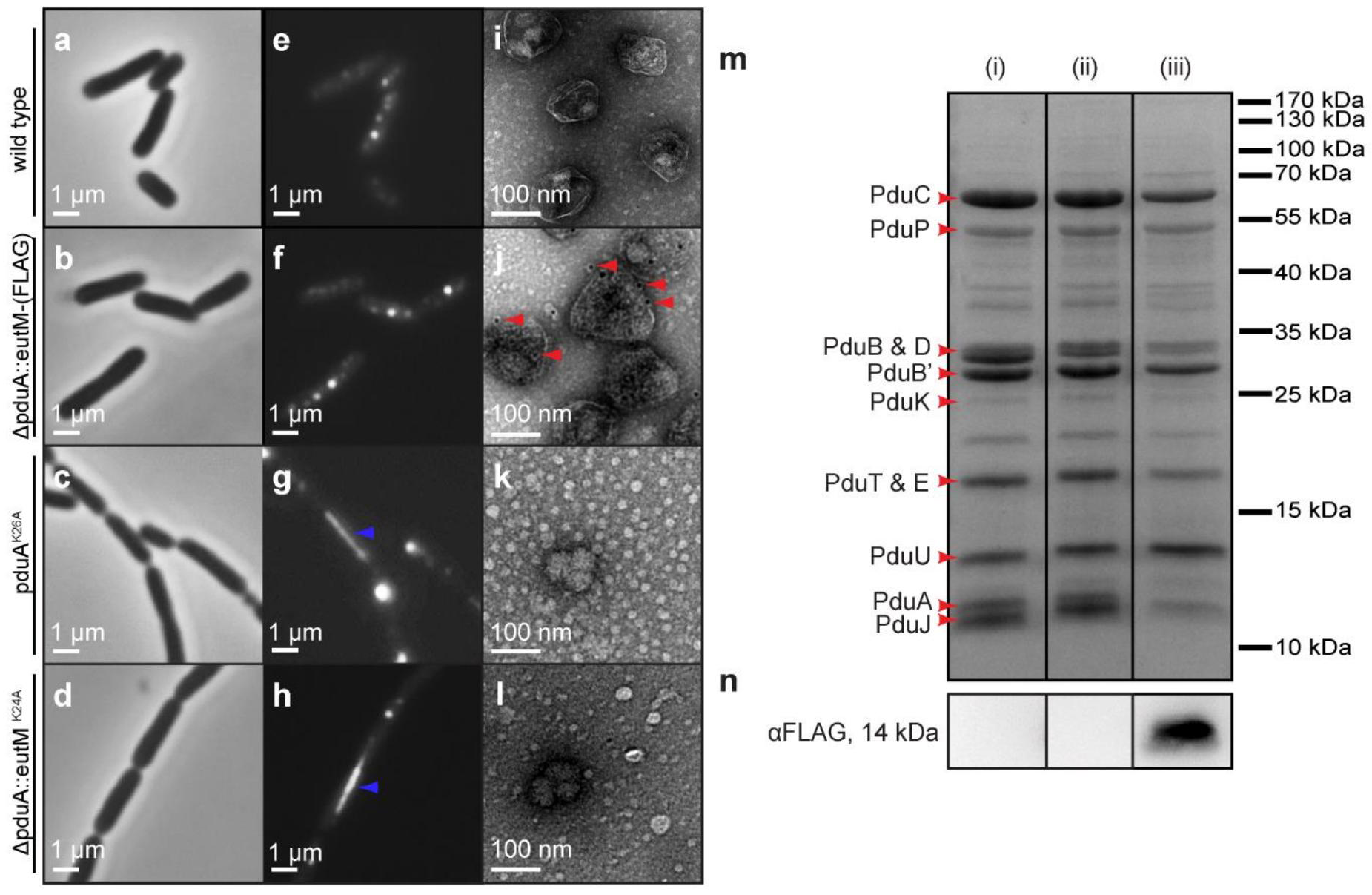
EutM incorporates into the Pdu MCP (a, b, c, d) Phase contrast images of *S. enterica* strains LT2 (*wild type*), LT2 *pduA: eutM*, LT2 *pduA*^*K26A*^, and LT2 *pduA: eutM*^*K24A*^, respectively. All strains are co-expressing Pdu MCPs and PduD^1-20^-GFP-ssrA. (e, f, g, h) GFP fluorescence microscopy images corresponding to phase contrast images a-d. Large filamentous aggregates are indicated with blue arrows in panels g and h. (i, j) Immunostained transmission electron microscopy of MCPs purified from wild type LT2 and LT2 *pduA: eutM-FLAG*, respectively. Bound IgG-conjugated gold particles are indicated with red arrows. (k, l) Transmission electron microscopy images of MCPs purified from the strains depicted in images c, d. For convenience, larger versions of the images in panels (i-l) are provided in Supplemental Figure S16. (m) Coomassie-stained SDS-PAGE of purified MCPs. Lane (i) is a purification from strain LT2 wild type, lane (ii) is from strain LT2 *pduA*^*K37Q*^, lane (iii) is from strain LT2 *pduA: eutM-FLAG*. (n)-FLAG western blot of purified compartments.

The protein content of purified MCP samples was analyzed by densitometry measurements of coomassie-stained SDS-PAGE gels. Prominent bands consistently observed in MCP samples were previously identified via mass spectrometry.^19^ Those identifications are used here. MCPs purified from the *pduA: eutM* strain exhibited shell protein bands in similar proportions to those found in wild-type MCPs, except that the PduA band was missing and the 13 kDa band expected to be PduU roughly doubled in density (Figures 2m, S4, S7). We hypothesized that this band represented both PduU and EutM superimposed. To verify the localization of EutM, the purification was repeated with a FLAG epitope tag fused to EutM. Western blotting indicated that EutM-FLAG runs at the apparent size of 13 kDa in a 12.5% acrylamide gel, and we confirmed that the 13 kDa band included EutM-FLAG by mass spectrometry (Figures 2n, S8, Table S1).

Next, we constructed a non-polar *pduA: pduA-FLAG* strain. We purified MCPs from this strain along with the *pduA: eutM-FLAG* strain and compared the levels of FLAG-tagged proteins using SDS-PAGE and western blot. Loading of the purified MCP samples was normalized by total protein concentration. We found that all samples contained similar amounts of PduA or EutM-FLAG (Figure S9).

To confirm that EutM is not merely co-purifying with Pdu MCPs, anti-FLAG immuno-gold antibodies were used to label Pdu MCPs purified from the *pduA: eutM-FLAG* strain. Immunostained TEM images show that the anti-FLAG immuno-gold antibodies bind well-formed MCP structures. In contrast, the wild-type MCP sample does not bind any gold particles (Figure 2 i, j). Taken together, all evidence supports our hypothesis that EutM properly incorporates into the Pdu MCP shell and causes no morphological defects.

### PduA^K26A^ and EutM^K24A^ mutations both abrogate MCP formation

Sinha *et al*. demonstrated that a PduA^K26A^ mutant does not form MCPs due to disruption of hexamer edge binding contacts, resulting in a growth defect on 1,2-PD even though the enzymes required to metabolize 1,2-PD are present.^27^ Also, MCP structures cannot be purified from the *pduA*
^
*K26A*
^ strain via established methods for purifying the Pdu MCP. We replicated these results (Figures 2k, S10, 3), and in addition used fluorescence microscopy with our encapsulation reporter PduD^1-20^-GFP-SsrA to observe this mutant *in vivo*. We did not observe an absence of fluorescence as we expected; instead, we saw cells linked by filaments resembling the nanotube or filament structures formed by overexpression of single 23, 27, 38 shell proteins or incomplete MCP operons as described previously (Figure 2c, g).^4, 28, 36, 37^ These structures were accompanied by bright puncta varying in appearance and predominantly found at the cell poles. These puncta are presumed to be aggregated or poorly formed MCPs since they cannot be purified in the same manner as non-mutated MCPs. Attempts to purify MCPs from this strain gave only dilute aggregate species as seen by SDS-PAGE analysis as well as by TEM (Figure 2k, Figure S10). The same phenotype was observed when a similar mutation was made to *eutM* in the *pduA: eutM* strain (Figure 2d, h, l, and Figure S10). Growth on 1,2-PD is decreased for a *pduA: eutM*
^
*K24A*
^ strain, likely due to mis-formed MCPs like those formed in the *pduA*
^
*K26A*
^ strain (Figure 3). *pduA: eutM*
^
*K24A*
^ and *pduA*
^
*K26A*
^ mutants were used as non-MCP forming controls in subsequent experiments. Their behaviors coincide in all tests. This is further evidence that EutM is able to complement PduA and fully integrate into the Pdu MCP. Moreover, our results indicate that the same lysine binding partner is important for assembly of both the wild-type Pdu MCP and the EutM Pdu MCP chimera.

**Figure 3.**
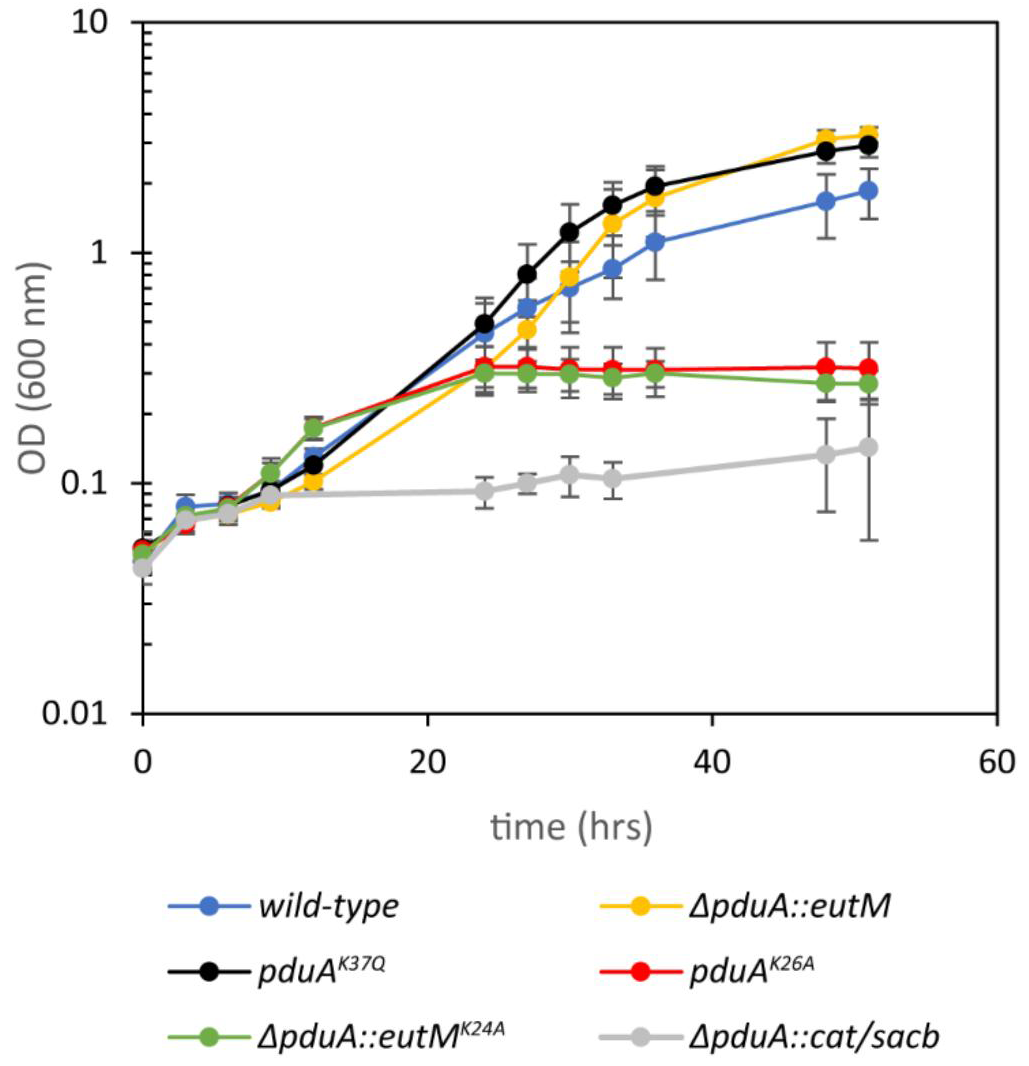
Strains harboring shell protein variants grow on 55 mM 1,2-PD and 150 nM coenzyme B12 MCP variant strains were grown on 55 mM 1,2-PD with 150 nM coenzyme B12. Wild-type strain LT2 is compared to LT2 *pduA: eutM*, *pduA*^*K37Q*^, *pduA*^*K26A*^, *pduA: eutM*^*K24A*^, and *pduA: cat/sacb* strains. Mean optical density at 600 nm is plotted vs time. Error bars represent 95% confidence interval from six measurements started on four different days.

### Limiting coenzyme B12 condition confirms*pduA: eutM* MCPs form a complete compartment shell

Coenzyme B12 is a required cofactor for the first step of 1,2-PD metabolism in the Pdu pathway. This cofactor is supplied in the growth medium because *S. enterica* cannot produce it under our aerobic conditions. Growth at low coenzyme B12 concentrations (20 nM) has been used previously as a method to sensitively detect whether mutations to MCP proteins result in non-assembly or defective MCP shells.^21,^

The limiting coenzyme B12 condition is advantageous for non-assembling MCP mutants, such as the *pduA*
^
*K26A*
^ mutant, and allows improved growth. Conversely, the wild type and mutant strains that assemble well grow poorly at 20 nM coenzyme B12 compared to mutants that do not assemble MCPs. We exploited this growth condition as an additional assay to detect if the mutations we made to PduA disrupted MCP assembly. We found that the *pduA: eutM* mutant grows as poorly as the wild-type strain in this condition (Figure S11), indicating that the chimeric MCP shells form a complete diffusion barrier.

### The effects of EutM substitution are recapitulated by a point mutation in the PduA pore

The above experiments showed that there were no observable differences in shell protein expression or MCP assembly between wild-type and *pduA: eutM* strains. Thus, we set out to investigate whether the behavior of the EutM-Pdu chimera is indeed due to a change in diffusion properties of the shell. We hypothesized that the difference in growth caused by the EutM substitution was the result of a change in diffusion through the central pore of the shell protein. To support this idea, we incorporated a chromosomal point mutation to encode PduA^K37Q^, to more closely mimic the EutM pore sequence.

The analyses performed for the *pduA: eutM* substitution mutant were repeated with *pduA*
^
*K37Q*
^. Strikingly, the *pduA*
^
*K37Q*
^ mutation confers improved growth (Figure 3, S2) over wild type while still maintaining normal MCP morphology and protein content (Figures 2m, S5, S6, S7, S9, S12, S13). A single point mutation to the pore causes the same effect as a complete shell protein substitution. This is strong evidence indicating that diffusion of molecules through the pore of the MCP shell is altered, causing the observed changes in growth and metabolite levels. A modification of lysine to glutamine changes the charge environment in the pore, and may also change the polarity or hydrogen bonding configuration. Both of these factors likely alter the diffusion barrier small molecules experience.^23, 39^ It is convenient for MCP engineering purposes that such small mutations to this region can confer improved function of the whole organelle structure.

### The EutM substitution and the PduA^K37Q^ pore mutant change the accumulation of downstream metabolites

HPLC analysis was performed to measure metabolite levels in the media over the course of growth on 1,2-PD (Figure 4). Fast growth on 1,2-PD has previously been associated with decreased aldehyde levels in the media.^10^ Therefore, we expected that the *pduA: eutM* and *pduA*
^
*K37Q*
^ strains would accumulate less aldehyde in the media. Indeed, we found that the accumulation of downstream products propionaldehyde, 1-propanol, and propionate are decreased in both the *pduA: eutM* and *pduA*
^
*K37Q*
^ mutants. We also found that the consumption of 1,2-PD occurred at roughly the same rate in wild-type, *pduA: eutM,* and *pduA*
^
*K37Q*
^ strains (the differences in calculated rate are not statistically significant).

**Figure 4.**
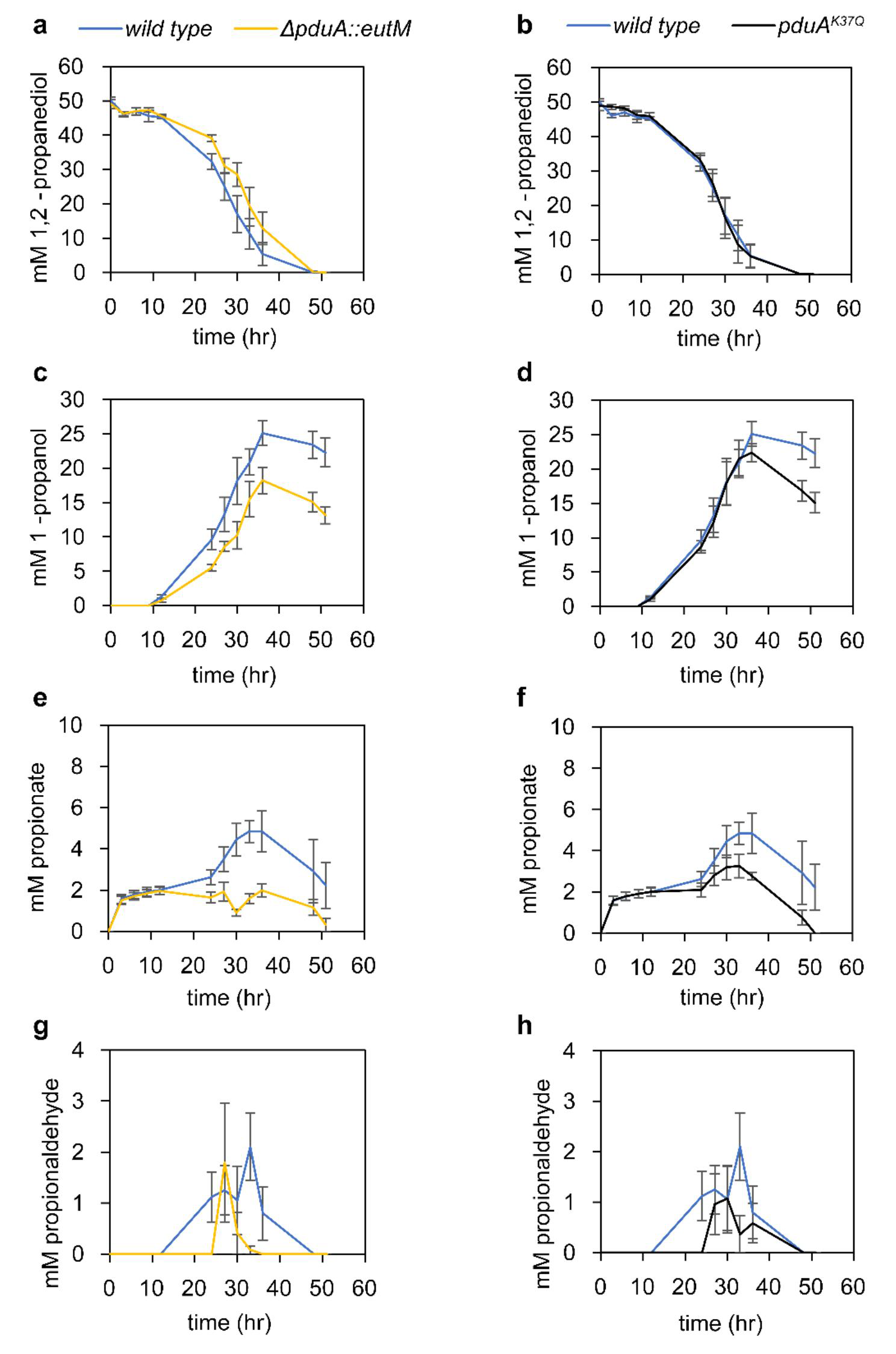
HPLC analysis of MCP-forming strains during growth on 1,2-PD and coenzyme B12 The concentrations of 1,2-PD and products of the Pdu pathway were monitored during growth on 55 mM 1,2-PD and 150 nM coenzyme B12. Mean values of metabolites are plotted vs time. Metabolites measured are 1,2-PD, 1-propanol, propionate, and propionaldehyde, as labeled. In the left hand column (a, c, e, g) the wild-type *S. enterica* LT2 is compared to LT2 *pduA: eutM*. In the right hand column (b, d, f, h) the wild-type strain is compared to LT2 *pduA*
^
*K37Q*
^. Error bars represent

Non-assembling mutants (*pduA*
^
*K26A*
^ and *pduA: eutM*
^
*K24A*
^) show the highest levels of propionaldehyde, at 8.6 and 10 mM, respectively, compared to 2.1 mM for the wild-type MCP (Figure 5). Propionaldehyde is toxic to *S. enterica* when exogenously added at concentrations of about 8 mM.^14^ Also, since propionaldehyde is volatile, accumulation of this molecule in the media results in some unmeasured amount of propionaldehyde evaporation to the head space. The differences in propionaldehyde concentrations among wild type, *pduA: eutM,* and *pduA*
^
*K37Q*
^ are more subtle. Wild-type *S. enterica* maintains about 2 mM propionaldehyde for much of the exponential phase. We observed no detectable propionaldehyde in *pduA: eutM* cultures apart from a single spike usually occurring between 27 and 30 hours reaching a level averaging 2 mM. The *pduA*
^
*K37Q*
^ strain behaved similarly to the *pduA: eutM* strain with respect to propionaldehyde accumulation. Therefore, while these strains reach similar maximum concentrations of propionaldehyde in the media, the *pduA: eutM,* and *pduA*
^
*K37Q*
^
*mutants*sustain this concentration for a shorter period.

**Figure 5.**
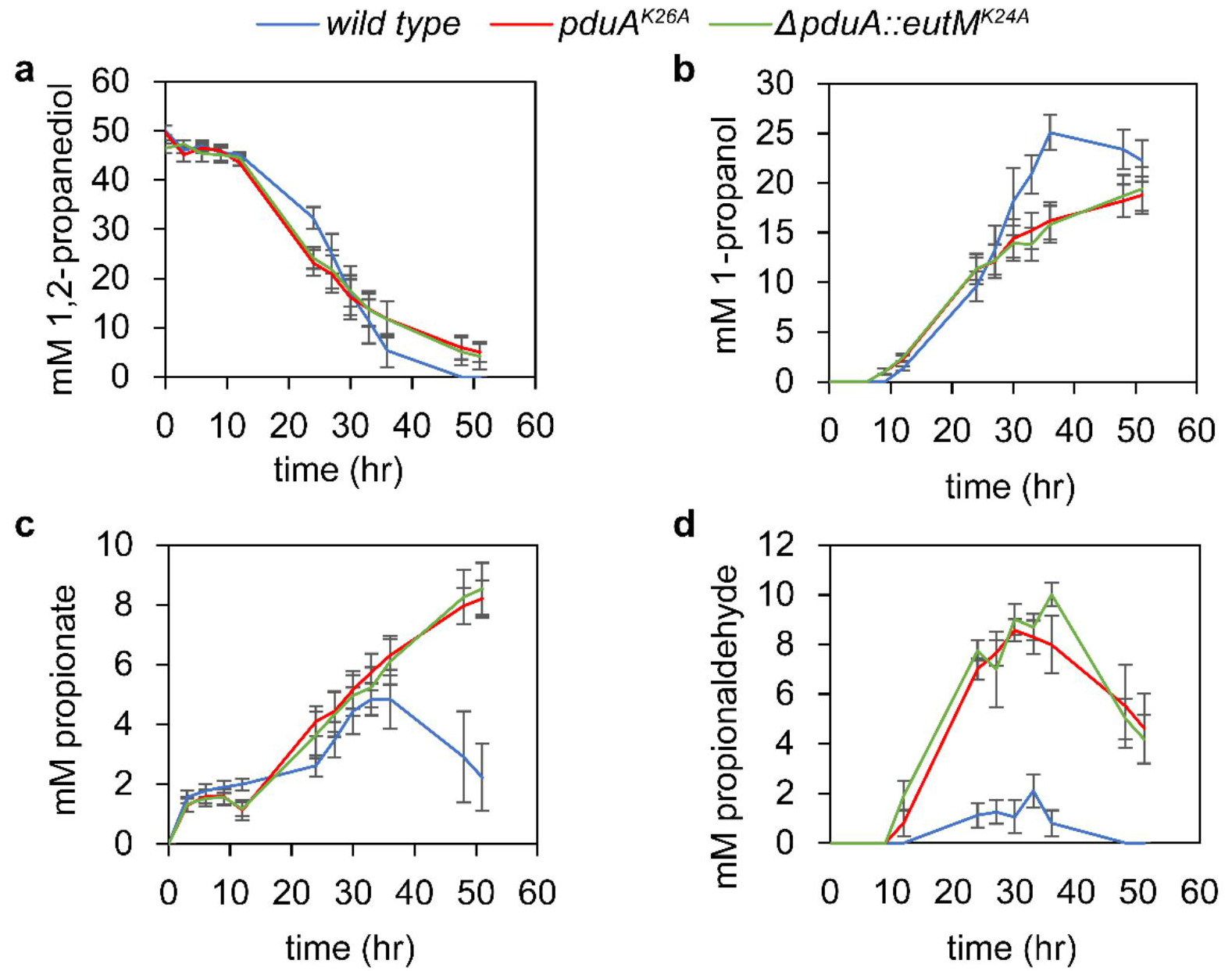
HPLC analysis of non-assembling shell protein mutant strains The concentrations of 1,2-PD and products of the Pdu pathway were monitored during growth on 55 mM 1,2-PD and 150 nM coenzyme B12. Mean values of metabolites are plotted vs time. Metabolites measured are 1,2-PD, 1-propanol, propionate, and propionaldehyde, as labeled. The wild-type LT2 strain is compared to LT2 *pduA*
^
*K26A*
^and LT2 *pduA: eutM*
^
*K24A*
^ strains. Error bars represent standard error from six measurements started on four different days.

In wild-type cultures the 1-propanol concentration level stabilizes at around 25 mM from 36 hours onward. In contrast, *pduA: eutM* and *pduA*
^
*K37Q*
^ strains reach a maximum of about 20 mM 1-propanol, and the concentration decreases after 36 hours. This indicates that the accumulation rate of 1-propanol is slower in the *pduA: eutM* and *pduA*
^
*K37Q*
^ mutants. Non-assembling mutants accumulate even less 1-propanol, likely due to the greatly increased escape of the aldehyde before conversion. After the supply of 1,2-PD is exhausted, 1-propanol in the media is likely converted back to propionaldehyde and then consumed.^23^ This process in addition to some evaporation from the headspace results in the decline in 1-propanol concentration at the end of each growth period. A surprising trend is seen in the concentration of propionate observed in the media over time. We observed increased propionate accumulation in strains that grew poorly. Non-assembling mutants ranked highest in propionate accumulation, followed by the wild type, then *pduA*
^
*K37Q*
^ and *pduA: eutM*.

To offer an explanation for these behaviors, we propose that the *ΔpduA: eutM* and *pduA*
^
*K37Q*
^ mutants restrict the diffusion of intermediate propionaldehyde, the negatively charged propionyl-phosphate product, or both. In the following discussion, we consider how this hypothesis is supported by the observed metabolite concentrations and growth phenotype.

The restriction of propionaldehyde diffusion by the *ΔpduA: eutM* and *pduA*
^
*K37Q*
^ mutants would directly result in the observed decreases in propionaldehyde accumulation compared to the wild-type. Since propionaldehyde is volatile, increased accumulation in the media leads not only to toxicity, but also to a loss of carbon to the gas phase. Preventing aldehyde escape may also result in a higher local propionaldehyde concentration and a kinetic enhancement of pathway throughput. Both of these effects contribute to the differences in final cell density observed between strains, leading to fastest growth in the *ΔpduA: eutM* and *pduA*
^
*K37Q*
^ mutants.

Restriction of propionyl-phosphate diffusion could result in the decreased accumulation of both propionate and 1-propanol by the *ΔpduA: eutM* and *pduA*
^
*K37Q*
^ mutants. To illustrate, non-assembling mutants allow free diffusion of propionyl-phosphate, facilitating its conversion to propionate by the PduW kinase, which is not localized to the MCP. In contrast, propionyl-phosphate molecules face a moderate diffusion barrier in the wild-type pore, resulting in decreased propionate release compared to non-assembling mutants. Thus, the *ΔpduA: eutM* and *pduA*
^
*K37Q*
^ mutants may form a greater diffusion barrier since even less propionate is released than in the wild-type strain. The mitigation of 1-propanol accumulation in *ΔpduA: eutM* and *pduA*
^
*K37Q*
^ mutants might also be due to restricted propionyl-phosphate diffusion. Decreased product escape could cause a pathway back-up reducing the rate of NADH production, leading to slower turnover in the 1-propanol branch of the pathway involved in NADH recycling. Considering pore structure, it is intuitive that a positively charged lysine in the wild-type PduA pore would facilitate the interaction of a negative product such as propionyl-phosphate more than the neutral glutamine, and allow a greater rate of egress.^39, 40^ Propionate escape in the wild-type could lead directly to decreased growth rate, or slower propionate uptake into central metabolism could be a result of aldehyde toxicity.

We recognize that there is still much to learn about the fate of propionate and the relationship between the 1,2-PD utilization pathway and the rest of central metabolism. According to our current understanding, the effect of decreased propionaldehyde leakage should result in flux enhancement. However, though strains that appear to restrict aldehyde leakage grow better, they also release less of the downstream products, propionate and 1-propanol. Further study is necessary resolve these two effects.

### Shell protein pore residues can be altered to control encapsulated enzyme function

We further explored the relationship between the pore and 1,2-PD metabolism by creating a genomically-encoded library of PduA variants that differ at residue 37. We checked each mutant for MCP formation by fluorescence microscopy using our encapsulation reporter PduD^1-20^-GFP-ssrA (Figure S12). All mutants exhibited fluorescent puncta, indicating the formation of MCPs. Growth on 1,2-PD with a limiting concentration of coenzyme B12 further demonstrated that all mutants did not form “leaky” MCP structures (Figure S11). Finally, we selected a few of the library members for purification. Purified MCPs from these strains are all similar to wild-type MCPs when examined by SDS-PAGE banding pattern and by TEM. (Figure S10 and S13).

The library was then screened for growth in NCE with 1,2-PD and 150 nM coenzyme B12 (Figure S14). The end-point cell density for each mutant is summarized in Figure 6a. To analyze these data we compared average end-point cell density to the characteristics of the amino acid substituted in each mutant. The variables considered were the predicted amino acid pKa, hydropathy index, and residue molecular weight. We found a moderate negative correlation between hydropathy index and final cell density (correlation coefficient of −0.47), indicating more hydrophobic side chains generally caused a decrease in growth (Figure 6c). Final cell density was also plotted versus pKa for several charged amino acid side chains. There was a negative correlation (correlation coefficient of −0.6) suggesting that a more positively charged pore is detrimental to growth on 1,2-PD (Figure 6b). There was no clear trend when growth was compared to size of the amino acid side chain alone (Figure S15). This indicates that, of the three characteristics considered, the hydrophobicity and charge of the amino acids in the pore were most important for retention of small molecules like propionaldehyde within the MCP.

**Figure 6.**
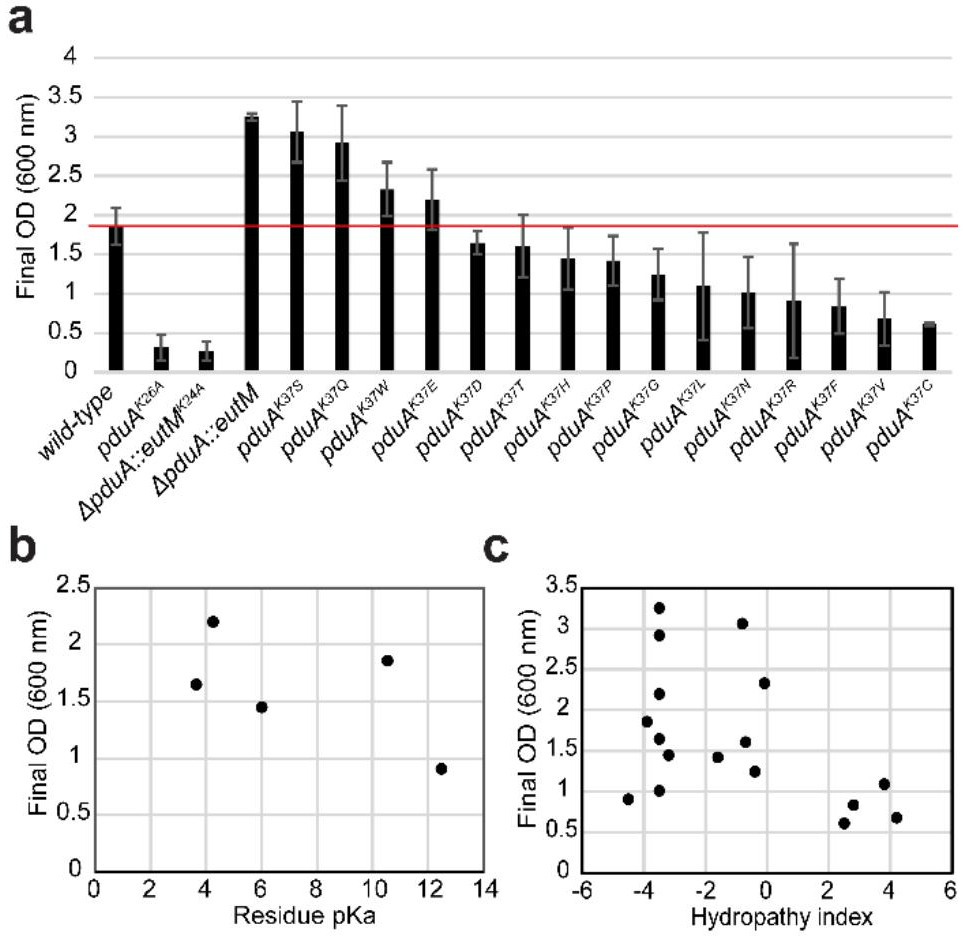
A library of amino acids at residue 37 of PduA confers a variety of growth effects (a) Mean values of final OD (600nm) are plotted for various PduA mutants from growth on 1,2-PD with 150 nm coenzyme B12. Error bars represent standard error from four different measurements started on separate days. The red line highlights wild-type growth for comparison. (b) Plot of final OD vs pKa of charged amino acids substituted at position 37 of PduA. (c) Plot of final OD vs hydropathy index of amino acids substituted at position 37 of PduA.

### Pore residues of PduA evolve rapidly

Upon analysis of the PduA K37 mutagenesis library, we were surprised to observe that the pore loop of PduA is very tolerant to mutation; indeed, all mutants formed MCPs similar to wild type. It is remarkable that this locus of PduA can accommodate any amino acid without a compensating mutation. For future engineering of the MCP shell, it is desirable to know if other residues may be mutated to change MCP function or to add chemical handles for the conjugation of useful moieties. We therefore used the Rate4Site program (version 2.01) to calculate an evolutionary rate scaling factor for each residue of PduA relative to the overall evolution rate of the protein.^41^ The input of this program is a multiple sequence alignment of 193 non-redundant PduA homologs from a variety of MCP systems. The resulting output is mapped over the structure of PduA (Figure 7a). Residues of a darker red color have high evolutionary rates, while white residues evolve slowly relative to the average rate for the entire protein. The evolutionary rate parameter represents the number of non-synonymous replacements expected per unit time for a specific residue. This parameter is related to conservation score but the rate calculation also uses the architecture and branch lengths of the phylogenetic tree in order to account for evolutionary time. If we detect residues that change often over time, but greatly affect the protein function, this could indicate a structural feature important for adapting function to environmental change.

**Figure 7.**
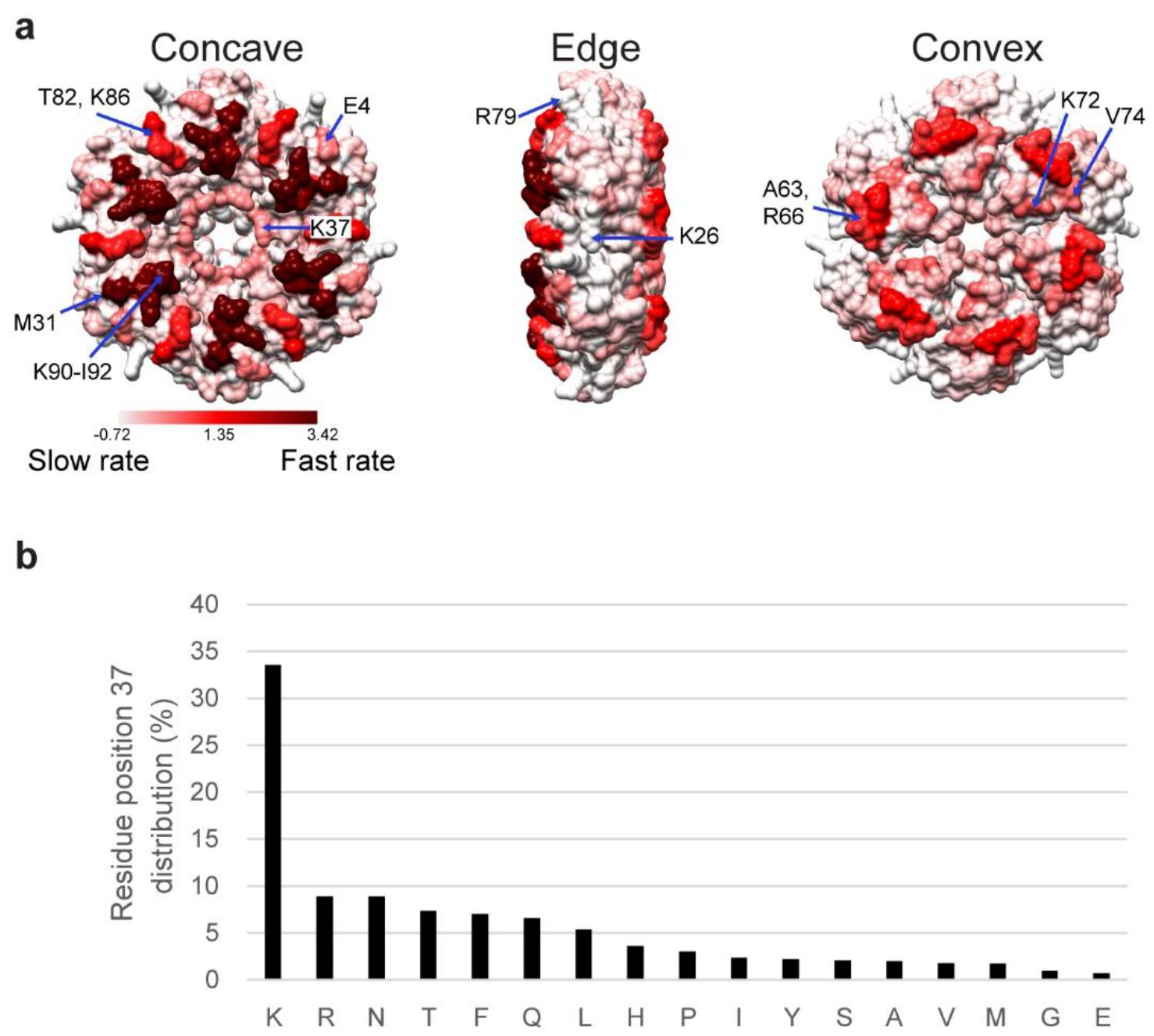
Phylogenetic analysis of PduA homologs (a) Rate4Site evolutionary rate scaling factor mapped over the crystal structure of PduA. White represents most conserved residues. Dark red represents residues with high evolution rates. Some residues with highest and lowest evolutionary rates are labeled for reference. (b) Histogram showing the % frequency of each amino acid aligning with PduA K37 in an MSA of n=1000 homolog sequences.

The results indicate that pore positions 37 and 40 have a higher evolutionary rate than the average evolution rate for the entire protein, while the isoleucine and glycine at positions 38 and 39 evolve more slowly than the protein average. This suggests that a hydrophobic motif at the narrowest part of the pore is essential for MCP function in general, whichever pathway is encapsulated within. On the other hand, while the residues forming rings on either side of the hydrophobic motif evolve more quickly, they evidently also affect encapsulated pathway function, most likely by inhibiting or facilitating diffusion of one or more metabolites. The mutability of certain pore residues may allow the flexibility to encapsulate a number of pathways involving a diversity of small molecules.

Interestingly, the residues K90-I92 near the C-terminus of PduA, as well as several other residues on both concave and convex surfaces, are also very rapidly evolving (labeled on Figure 7a). These areas of the protein structure are less constrained and may serve as good targets for protein modifications, particularly for control of cargo loading or for labeling the outer surface for engineering applications.

### PduA pore mutants conferring faster growth on 1,2-PD are not abundant among PduA homologs

Among the strains in the K37 mutation library, the strains with Gln, Glu, Ser, and Trp at position 37 grew to a greater final OD600 than the wild-type strain under laboratory conditions. Curiously, this indicates that the *S. enterica* serovar lab strain used in this study is not optimized for flux through the Pdu pathway and other factors may affect the fitness of LT2 in the host gut. We searched among PduA homologs to find out if any of the amino acid variants that resulted in highest growth were among those commonly found in nature. A distribution of the frequencies of each amino acid aligned with position 37 is given in Figure 7b. Together, strains with Gln, Glu, Ser, or Trp aligned with position 37 only represent 9.38% of over 1000 related sequences. The population of PduA homologs in nature is not enriched for these high performers.

## Conclusions

In this work, we demonstrate the ability to assemble Pdu MCPs with a foreign shell protein substitution, and the flexibility to use protein engineering to alter the pore structures of PduA without abrogating MCP structure. For the first time, mutations to the Pdu MCP shell have resulted in enhanced function of the enclosed pathway. PduA shell protein mutants, including those with only a single amino acid substitution of a pore-lining residue, have improved growth phenotypes with 1,2-PD as a carbon source. We uncovered significant evidence that these phenotypes are due to modification of small molecule diffusion rates through the pores of the MCP, leading to escape of the intermediate aldehyde and possibly preventing the escape of the product propionyl-phosphate. These trends are observable through HPLC analysis of metabolites and are supported by our analysis of protein structure and MCP assembly. An investigation of the relationship between amino acid chemistry in the pore and the resulting growth performance indicates that charge and hydrophobicity of the side chain play key roles. Shells with more negatively charged pores tended to grow poorly on 1,2-PD, while an increase in the hydropathy of the residue at position 37 also decreased the fitness of the strain. The PduA K37 mutagenesis library produced a range of growth phenotypes with strains that grew on 1,2-PD faster or slower than the wild type. Generation of chimeras and/or modification of shell protein pore residues are both promising techniques to optimize MCP shell characteristics for an encapsulated pathway of interest and to achieve useful phenotypes not found in nature.

## METHODS

### Bacterial strains, media, and growth conditions

The bacterial strains used in this study are *Salmonella enterica* serovar Typhimurium LT2 and *Escherichia coli* DH10B. For mutations to *pduA*, the gene was amplified from the *S. enterica* chromosome and inserted into a pBAD33, p15a, cm^R^ plasmid by golden gate assembly with BsaI restriction endonuclease.^42^ This plasmid was transformed into *E. coli*. *pduA* and was mutated as described above using PCR site-directed mutagenesis. EutM K24A was created in the same manner. Primers for each mutation are shown in supplemental Table S2.

Chromosomal modifications were made by recombineering as described by Court.^43^ First, the native *pduA* gene was replaced with the *cat/sacb* cassette amplified from the TUC01 genome with primers mfsp105 and either mfsp106 or mfsp195 for *eutM* or *pduA* insertions, respectively (primer sequences listed in Table S2). This amplicon was used to create *pduA: cat/sacb*. The e*utM* gene was amplified with primers mfsp199 and mfsp200 and used to make *pduA: eutM*. To make mutations in *pduA,* the gene was first subcloned into a plasmid vector and mutated by site directed mutagenesis as described above. The inserts were then amplified from plasmids by PCR with primers mfsp197 and mfsp104 and used to make all chromosomal *pduA* mutants. For FLAG tag fusions, the non-polar *pduA: cat/sacb* strain constructed with mfsp106 was used as a starting point to maintain the same RBS region for the subsequent *pduB* gene. The FLAG sequence was added by PCR to *pduA* and *eutM* sequences using primers mfsp304 and mfsp218, respectively before insertion by recombineering. Each strain was confirmed by Sanger sequencing of PCR amplicons from the PduA region of the *S. enterica* chromosome.

For growth of *S. enterica*, single colonies were picked from freshly streaked plates and were grown in 5 mL of TB (Terrific Broth) medium at 30°C, 225 rpm, for 24 hours. For growth curves with a 1,2-PD carbon source, cultures were diluted to an OD600 of 0.05 in 50 mL No-Carbon-E (NCE) minimal medium supplemented with (unless stated otherwise) 55 mM 1,2-PD, 150 nM coenzyme B12, 1 mM magnesium sulfate, and 50 μM ferric citrate.^32^ Cultures were grown in 250 mL flasks at 37°C in an orbital shaker at 225 rpm. At every time point, 1.5 mL of cell culture sample was taken for OD600 measurement and HPLC analysis. The doubling time of each growth curve was calculated by plotting the log of the OD600 vs. time. The linear region of each individual curve was identified (including a minimum of three time points). The slope of the linear regression between time points was determined, and the doubling time was calculated with = log(2) /. For MCP purifications, flow cytometry, or microscopy experiments, single colonies were grown in 5 mL LB (Lysogeny broth)-Lennox medium at 30°C, 225 rpm, for 24 hours. Cell cultures were diluted from the primary culture 1:1000 into NCE supplemented with 42 mM sodium succinate, 1mM magnesium sulfate, and 50 μM ferric citrate. For cultures containing the plasmid expressing PduD^1-20^-GFP-ssrA, half the usual amount of antibiotic was used (17 mg/mL chloramphenicol). Culture volumes were 400 mL of media in 2L flasks for MCP purification, and 5 mL of media in 24-well blocks (Analytical Sales and Services, Inc., cat. no. 24108) for flow cytometry and microscopy. Cells were grown at 37˚C in an orbital shaker at 225 rpm. For cultures with PduD^1-20^-GFP-ssrA expressed from a pBAD33 plasmid, 1.33 mM arabinose was added when subculture reached an OD600 of 0.4. After five additional hours of growth, samples were taken for fluorescence microscopy, flow cytometry, or Pdu MCP purification.

### High Performance Liquid Chromatography (HPLC) media analysis

At each time point, a 750 µL sample of whole cell culture was taken. This was centrifuged for 5 min at 13,000 g to remove cells. The supernatant was decanted and frozen until time of analysis. Before analysis samples were thawed and filtered. The Shimadzu HPLC detection system was equipped with a Bio-Rad Aminex HPX-87H (300 by 7.8 mm) ion exclusion column, LC-20AD solvent delivery system, SIL-20AC auto sampler, RID-10A refractive index detector, and SPD-M20A diode array detector (Shimadzu, Kyoto, Japan). Mobile phase is 5 mM H2SO4 with 0.4 mL min^-1^ flow rate.^14^ The column temperature is 35°C.

### MCP purification

MCPs were purified by centrifugation of lysate as described previously.^44^ They were stored at 4°C until analysis by SDS-PAGE or TEM. Total protein concentration of pure MCP sample was determined by Pierce BCA protein assay kit (Thermo Fisher).

### Transmission electron microscopy

Samples were placed on 400 mesh formvar coated copper grids with a carbon film after the grids were treated by glow discharge. 10 μL of purified MCPs at a concentration of about 0.1 mg/mL was placed on each grid for 2 min. The grids were washed three times with deionized water before fixation. Then, 10 μL of 2% glutaraldehyde in water was placed on each grid for 1 min. Grids were then washed an additional three times with deionized water. Finally, samples were stained in 1.6% aqueous uranyl acetate for 1 min. All solutions were centrifuged directly before use to avoid aggregate or particulate contact with grids. Samples were imaged with an REI Tecnai T12 transmission electron microscope and a Gatan Ultrascan 1000 camera (Gatan, Pleasanton, CA).

### Immunostaining

Samples were placed on 400 mesh formvar coated copper grids with a carbon film after glow discharge treatment. 10 L of purified MCPs at a concentration of about 0.1 mg/mL was placed on grids for 2 min. The grids were washed three times with deionized water. Blocking buffer was prepared with 25 mL of phosphate buffered salt (PBS) at pH 7.4, 0.2 g bovine serum albumin (BSA), and 25 μL of gelatin from cold water fish skin (Sigma). Then, grids were incubated with blocking buffer for 15 min. Grids were incubated with mouse anti-FLAG primary antibody (Sigma) diluted 1:1000 in the blocking buffer for 2 hours. Grids were washed four times with PBS for 2 min each. Grids were then incubated with anti-mouse IgG gold secondary antibody produced in goat (Sigma) diluted 1:20 in blocking buffer for 1 hour. Grids were washed again four times with PBS for 2 min each time. Samples were fixed with 2% glutaraldehyde in PBS for 5 min. Samples were washed for 2 min in PBS, then four times in deionized water. Grids were stained with 1% uranyl acetate in water for 3 min before grid was allowed to dry.

### Mass spectroscopy

Protein bands were excised manually and transferred to Eppendorf tubes (0.2 mL). Protein-containing gel pieces were washed with a series of washes that included alternating additions of 100 μL 0.1 M ammonium bicarbonate followed by 100 μL of 100% ACN in order to remove traces of coomassie staining. Once gel slices were thoroughly destained, they were dried using a speed-vac. Gel pieces were then subjected to 10 mM dithiothreitol for 20 minutes at 56 °C and subsequently alkylated with 20 mM iodoacetamide for 1 h at room temperature in the dark. Acetonitrile was then added to shrink the gel pieces and the pieces were thoroughly dried in the speed-vac prior to addition of 0.1 ml of a solution containing 12.5 ng/ l trypsin in 50 mM ammonium bicarbonate. Gel pieces were allowed to swell at 4C for 3 hours prior to overnight digestion at 37 °C. Liquid was collected following digestion and the gel pieces were bathed in 100% acetonitrile to adequately shrink the gel pieces. All liquid was collected into one tube and evaporated to dryness using a speed-vac. The peptide solution was reconstituted into 10 μl of 2% ACN with 0.1% TFA and assessed for peptide concentration based on the absorbance at 280 nm using a Nanodrop spectrophotometer. One microgram total peptide was loaded onto the nano-ACQUITY UPLC™ chromatographic system. Peptides were loaded and separated on a C18 Trizaic Naotile using a 60 min RP gradient at 450 nL/min (3–40% ACN over 40 min). The column temperature was set at 45 °C. Lock mass (Leucine enkephalin (556.2771 Da), 250 fmol/mL) was constantly infused by the NanoAcquity auxiliary pump at a constant flow rate of 1 μL/min with lockspray scans set at intervals of 45 seconds. The Xevo QTof ™ mass spectrometer (Waters) was programmed to switch between low (6 eV) and high (18–42 eV) energies in the collision cell, with a scan time of 1 s per function over a mass range of 50–2000 Da. LC-MS^E^ data were processed with ProteinLynx GlobalServer v2.3 (Waters) and searched in the associated *S. enterica* protein database (UniProtKB/SwissProt Protein Knowledge Base).

### Protein electrophoresis and western blot

Denaturing protein electrophorese (SDS-PAGE) was performed on purified MCPs using a 12.5% polyacrylamide gel unless otherwise noted. A 130V potential was applied for 70 minutes. Unless otherwise noted, equal volumes (15 µL) of each MCP sample were loaded onto the gel, and densitometry quantifications were normalized by sample protein concentration. Gels were either stained with Coomassie dye or further processed for western blot.^44^ Western blotting was done with a PVDF membrane according to standard protocols. Blotting was done with a mouse anti-FLAG primary antibody (Sigma) 1:2000 dilution in 50 mM Tris 150 mM NaCl pH 7.6 with 0.05% Tween-20 (TBST) with 1% w/v dry milk. The secondary antibody was HRP-conjugated goat anti-mouse antibody (Thermo) diluted 1:1000 in TBST. Labeling was visualized with west-pico chemiluminescent substrate (Thermo) using a Bio-Rad ChemiDoc XRS+. Densitometry analysis of Coomassie stained gels was performed using at least three different MCP purifications for each strain. Each purified sample was run on three different gels. Images were analyzed using the Image Lab software (Bio-Rad). The absolute signal measurement for each band was normalized by the signal for the band containing proteins PduT and PduE (band 7 in Figure 2). This band had the least variation in signal over all replicates.

### Fluorescence microscopy

Bacteria were viewed using a Nikon Ni-U upright microscope with a 100x 1.45 n.a. plan apochromat objective. Images were captured using an Andor Clara-Lite digital camera and Nikon NIS Elements software. Fluorescence images were collected using a C-FL Endow GFP HYQ bandpass filter. All images were taken with 400 ms exposure and adjusted identically for contrast in Adobe Photoshop software.

### Flow cytometry

Cultures were grown as described above. Samples were diluted 1:40 in phosphate-buffered saline (PBS) supplemented with 2 g/L kanamycin (to halt translation) in 96-well plates. 10,000 events were collected for each sample on a Millipore Guava easyCyte 5HT instrument. Cells were distinguished from debris by gating on the forward and side scatter channels using the FlowJo software. Reported fluorescence values are the arithmetic mean of the geometric mean green fluorescence of three independent samples acquired on three different days. Error bars represent one standard deviation.

### Multiple Sequence Alignment construction and evolutionary rate analysis

Evolutionary rate of each amino acid locus of PduA was calculated using the Rate4Site program version 2.01.^41^ The input for this tool was a multiple sequence alignment (MSA) of 193 non-redundant PduA homologs. The homologs were retrieved and the MSA assembled by the jackhmmer web server (https://www.ebi.ac.uk/Tools/hmmer/search/jackhmmer). Rate4Site uses this data to construct a phylogenetic tree. With both MSA and tree information, a rate-scaling factor is calculated that indicates how rapidly each residue evolves relative to the mean protein rate. Default settings (including Bayesian framework) were used here. A larger MSA of about 1000 sequences, also retrieved and aligned with jackhammer, was used to calculate amino acid frequency.

## SUPPORTING INFORMATION

Supplementary information including tables and figures cited in the text is available free of charge via the Internet at http://pubs.acs.org.

## ABBREVIATIONS

MCP: microcompartment 1,2-PD: 1,2-propanediol
GFP: green fluorescent protein
Pdu: 1,2-propanediol utilization
Eut: ethanolamine utilization
NCE medium: No-Carbon-E medium
TEM: transmission electron microscopy
MSA: multiple sequence alignment
PBS: phosphate buffered salt

## Author Contributions

M. F. S and D. T. E. designed the project. M. F. S. performed the experiments. M. F. S, C. M. J, and D. T. E. wrote the manuscript.

## Notes

The authors declare no competing financial interest.

## ACKNOWLEDGEMENTS

The authors wish to thank Chuchu Zhang for development of the Cell profiler pipeline used in this work, as well as the Robert D. Ogg Electron Microscope Laboratory for equipment usage, training, and materials. The authors would also like to thank the Tullman-Ercek lab for thoughtful discussions and comments. The work was funded by the National Science Foundation (award MCB1150567 to D. T. E.).

